# Effect of AT1 receptor blockade on cardiovascular outcome after cardiorespiratory arrest: an experimental study in rats

**DOI:** 10.1101/2023.05.16.540941

**Authors:** EAF Araújo Filho, MJC Carmona, DA Otsuki, DRR Maia, LGCA Lima, MF Vane

## Abstract

**Background:** AT1 angiotensin II receptor (ATI) antagonists are beneficial in focal ischemia/reperfusion (I/R) cases. However, in cases of global I/R, such as cardiorespiratory arrest (CRA), ATI blocker’s effects are still unknown.

**Methods:** Rats were allocated into four groups: Sham group (SG) – animals submitted to surgical interventions, without CRA; Control group (CG) – animals submitted to CRA and ventricular fibrillation; Group AT1 (GAT1) – like CG, plus 0.2 mg/kg of Candesartan; Vehicle Group (VG): animals equally induced to CRA, and administration of 0.2 ml/kg of dimethyl sulfoxide. The rate of return of spontaneous circulation (ROSC), survival, hemodynamic variables, histopathology, and markers of tissue injury were analyzed.

**Results:** Compared to CG, the GAT1 group had a higher rate of ROSC (62.5% vs. 42.1%, p<0.0001), survival (100% vs. 62.5%, CI: 0.014-0.034; p = 0.027), lower incidence of arrhythmia after 10 minutes of ROSC, (10% vs. 62.5%, p=0.000) and lower neuronal and cardiac injury scores (p=0.025 and p=0.021, respectively). The groups did not differ regarding CRA duration, number of adrenaline doses, or number of defibrillations.

**Conclusion:** ATI receptor blockade was responsible for higher rates of ROSC and survival, in addition to demonstrating neuronal and myocardial protection.

**Highlights:**

1. AT1 receptor block was responsible per higher rates of ROSC and survival in intervention group.
2. The AT1 receptor block can be neuroprotector in ischemic injury caused by CPR.
3. The candesartan administration during CPR can contribute with reduction of ventricular arrythmias.

## Introduction

Cardiorespiratory arrest (CRA) is the most severe adverse event in the perioperative scenario. Even if promptly assisted, it has a high mortality rate, ranging from 35% immediately after the event to approximately 70% after one year ^1–4^. Several studies have discussed ways to intervene and reduce the damage caused by global ischemia due to CRA ^5–7^.

The renin-angiotensin system (RAS), and its main effector, angiotensin II (Ang II), play a fundamental role in mediating oxidative stress, in addition to inducing endothelial dysfunction and activating pro-inflammatory pathways ^8^. Recent studies demonstrate using RAS inhibitors to reduce tissue damage in ischemia-reperfusion (I/R) ^8–11^. Angiotensin receptor antagonists (ARA) decrease areas of necrosis both in myocardial and cerebral I/R models ^12–14^. It is also suggested that using a specific ARA for the type 1 angiotensin II receptor (AT1), such as candesartan, Ang-II maintains effectiveness on type 2 receptors (AT2). The benefits of neuronal I/R models involve neuroprotection, decreased oxidative stress, and decreased edema after an ischemic event ^11, 14^.

There are no studies correlating the use of RAS blockers with the ischemia of multiple organs in CRA. The number of ischemic tissues is high during CRA events, leading to more aggressive I/R injuries. Thus, there is little information on whether blocking the RAS would be beneficial in this context.

Therefore, we plan to evaluate the role of angiotensin II AT1-receptor blockers during CRA. For that, an experimental study was conducted with Wistar rats submitted to CRA induced by ventricular fibrillation. The effects of RAS blockage were analyzed through intravenous administration of candesartan at the beginning of CRA maneuvers.

## Methods

An experimental study was conducted with isogenic male Wistar rats weighing 393g (380g – 470g), from the Central Animal Facility of the Faculty of Medicine of the University of São Paulo. The Ethics Committee approved the study for the Use of Animals of the University of São Paulo – CEUA-USP, protocol number: 034/16.

After being anesthetized and subjected to surgical procedures, the rats were allocated in a simple random fashion into four different groups.

Sham Group (SH): Anesthetized animals submitted to surgical instrumentation procedures for hemodynamic monitoring but with no ventricular fibrillation (VF) induction.

Control Group (CG): Animals submitted to CRA within 6 minutes of anoxia, followed by cardiopulmonary resuscitation (CPR). At the start of CPR, 20 mcg/kg of adrenaline and 0.5 ml of 0.9% saline were administered.

Vehicle group (VG): Animals submitted to CRA within 6 minutes of anoxia, followed by CPR. Adrenaline (20 mcg/kg) and 0.2ml/kg of 99% dimethyl sulfoxide (DMSO) were administered at the beginning of CPR.

Group AT1 (GAT1): Animals submitted to CRA within 6 minutes of anoxia, followed by CPR. At the beginning of CPR, adrenaline (20 mcg/kg), together with an intravenous dose of the candesartan cilexetil metabolite, Candesartan (>98%, CAS 139481-59-7), diluted in DMSO (99%) in 0.1%, at a dose of 0.2 mg/kg was injected.

### Instrumentation Procedures

After induction of general anesthesia with isoflurane (1 CAM), electrocardiographic monitoring and orotracheal intubation were performed with a 16 G flexible venous catheter (B. Braun). Next, the animals were placed on mechanical ventilation with a tidal volume of 8 ml/kg and a respiratory rate of 60 breaths per minute (bpm). A rectal thermometer adjusted the body temperature to 37 +/- 1oC. An incision was made in the cervical region, locating the right internal jugular vein. Then, a venous catheter was inserted for drug infusion with conduit for CPR induction. Next, the inguinal region was incised and another PE10 catheter was introduced into the right common femoral artery to collect parameters and invasively measure blood pressure using a data acquisition system (MP-100, BIOPAC, USA).

The electrocardiographic analysis was performed using a module coupled to the same system. CPR was induced by inserting a single lumen 3Fr catheter inserted through the jugular and guided by the pressure curve into the right ventricle, an electrode was introduced, and an electrical discharge of 1mA (9V, 60Hz) was applied for 3 minutes. Simultaneously, mechanical ventilation was discontinued. Then, the guide was removed, and the venous catheter retracted 1 cm and fixed for drug administration. The animals were suppressed for 3 minutes in CPR, totaling 6 minutes from the beginning of VF induction. Afterwards, cardiopulmonary resuscitation maneuvers were started with external chest compressions performed by an automatic compressor at a rate of 200 compressions per minute and depth of 1.5 cm 15. Ventilation (RR-25ipm) was reestablished with the administration of adrenaline (20mcg/kg) and sodium bicarbonate (1 mEq/kg). Heart rate was checked every 3 minutes. If VF was present, a 7J discharge was performed. CPR was restarted for another 3 minutes after defibrillation, with a new dose of adrenaline. After a heart rate check, arterial pulsations were taken as return of spontaneous circulation (ROSC). Maintenance of mean arterial pressure above 25 mmHg for 10 minutes was considered sustained ROSC and counted as mortality. If there were no pulsations in the second check and still in VF, a new 7J defibrillation charge was applied and chest compressions were resumed, adding a new dose of adrenaline. New defibrillations and rhythm checks were performed every 2 minutes, and adrenaline was administered every 3 minutes. After 20 minutes of CPR, efforts were terminated. However, if sustained ROSC and hemodynamic stability were established, the animals were observed for an additional 4 hours (Figure 1).

**Fig. 1.**
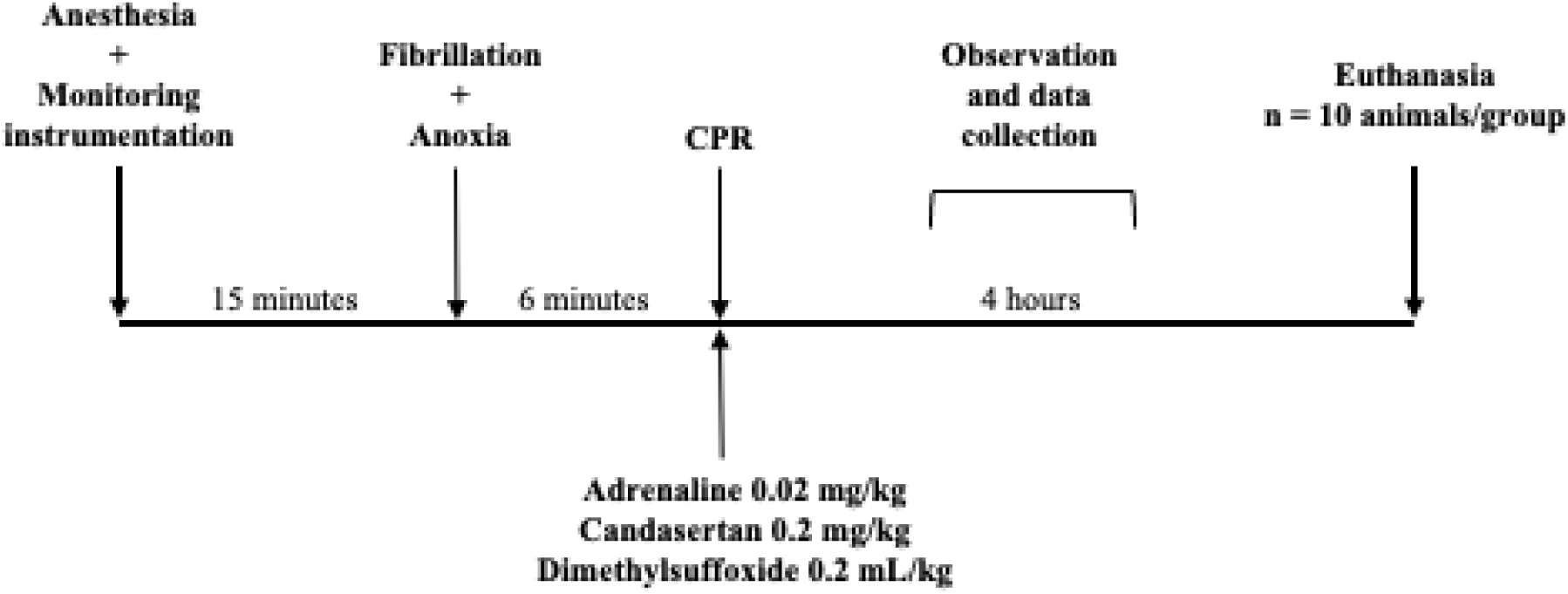
Experiment flowchart. CPR: cardiopulmonary resuscitation

The animals surviving the 4-hour period had a blood sample collected for troponin measurement. After, they were sacrificed by an overdose of inhalational anesthetic (6% isoflurane). Afterward, the heart and brain were extracted for processing and stored partly in formalin and partially frozen at -70°C.

#### Outcomes assessment

##### Hemodynamic evaluation

Blood pressure, heart rate, and cardiac rhythm were continuously recorded from the beginning of surgical instrumentation procedures, after anesthetic induction, and until animal sacrifice. The parameters were recorded on data sheets every 10 minutes, from the ROSC until the end of the experiment. Cardiac arrhythmias were classified according to the Lambeth convention guideline ^16^.

##### Gasometric determination

Blood samples (1mL in a heparinized syringe) were collected at pre-CPR induction, after CPR, and after 4 hours of observation. Samples were allocated in specific tubes and analyzed by an ABL800 Flex, Radiometer device.

##### Histology

After the animals were sacrificed, the brain and heart were carefully dissected and immediately fixed in 4% paraformaldehyde in phosphate buffer pH 7.0 for 24 hours. After, they were dehydrated in an alcoholic gradient (70° to 100°), cleared in xylol, and embedded in paraffin. 5µm cuts were obtained for morphological and morphometric evaluation.

The pathologist responsible for the analysis was blind to the animal group and the corresponding slide. The histopathological evaluation of both the brain and the myocardium was scored based on the percentage of lesions through manual quantification for each photomicrograph: 1= (<30%); score 2 = (31%–60%); score 3 = (>60%) ^18^.

##### TUNEL

The Terminal deoxynucleotidyl transferase-mediated (dUTP) nick end Labeling kit, also called the Tunel labeling index, was used to analyze cell injury and apoptosis. The slides were manually counted at 100x magnification. Twenty-five brain fields and 50 heart fields were analyzed. For the heart, 25 were from the right ventricle, and 25 were from the left. Means for each slide and the median for each group were then calculated. The slides were blindly read for each group.

##### Statistical analysis

All continuous variables are presented as Mean±SD after the Kolmogorov‒Smirnov test confirms the normality. Data were analyzed by one-way analysis of variance (1-way ANOVA) or 2-way ANOVA, where appropriate, followed by post hoc Tukey tests for comparisons between different groups. Non-normal data were analyzed by Kruskal-Wallis. Fisher’s exact test (chi-square) was used to analyze CPR rate, survival, and myocardial and brain injury scores. Statistical significance was assumed for P<0.05. Statistical analyzes were performed in SPSS version 23.0 for Windows (SPSS, Chicago, IL).

## Results

The study was conducted with 93 rats divided into four groups: 10 in the intervention, control, and vehicle groups, and 4 in the Sham group. Nine animals were discarded (five due to iatrogenic injury and four due to failure to induce CPR). Thus, 84 rats were included in the study. Among the 26 animals used in the vehicle group, only 3 met CPR criteria, and none completed the observational period. Thus, the group was excluded from the analysis (Figure 2).

**Fig. 2.**
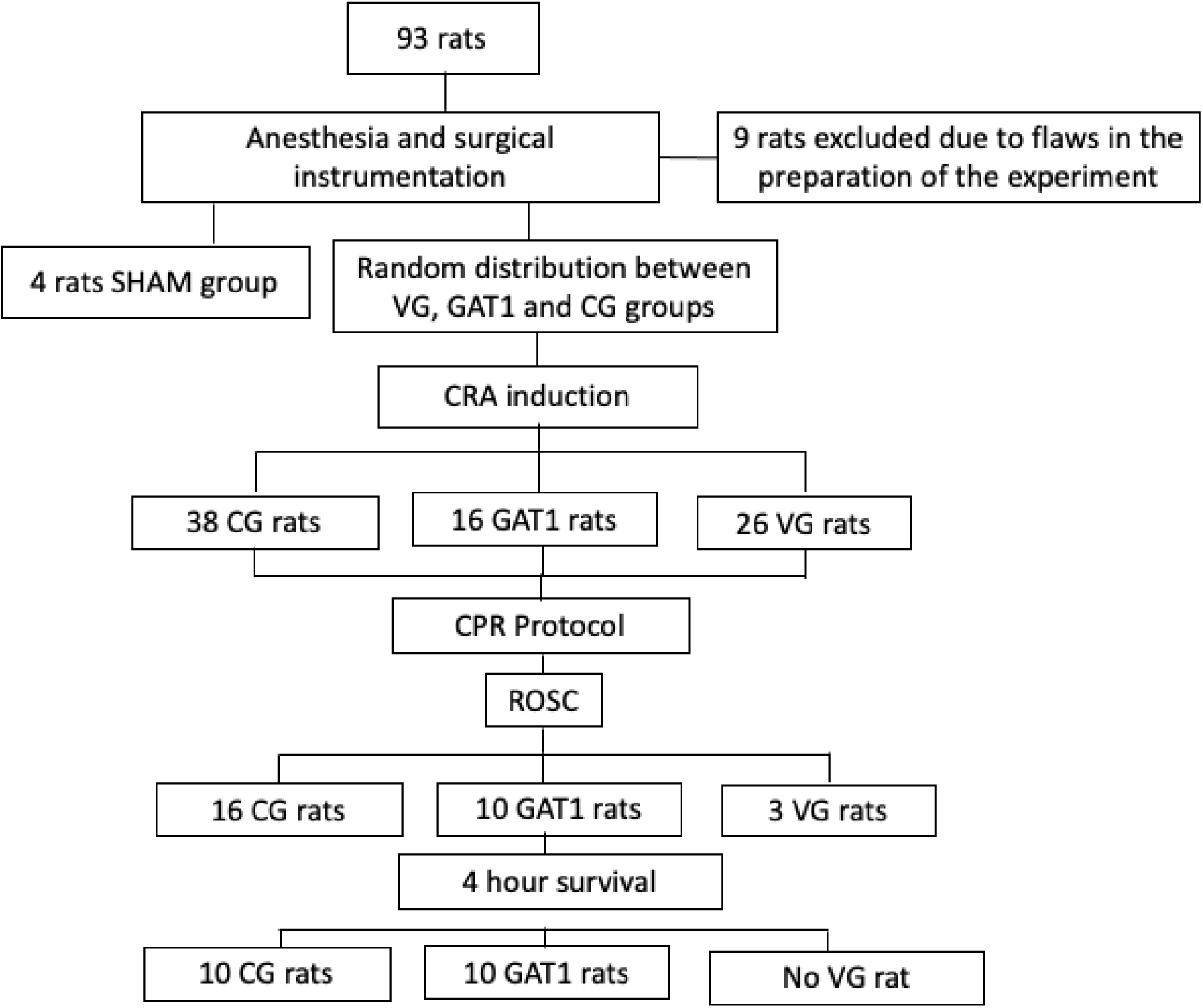
Distribution of rats in the study groups according to survival after cardiopulmonary resuscitation (CPR). CG: Control group; GAT1: Candesartan group; VG: Vehicle group; CPR: Cardiopulmonary resuscitation; ROSC: Return of Spontaneous Circulation.

### Baseline physiological parameters and survival

The average weight of the animals was: CG group 379.5 ± 47.4g, GAT1 group 409.7 ± 28.7g, and Sham group 465.4 ± 38.5g. The three groups had no significant differences in physiological and hemodynamic parameters (Table 1).

**Table 1.**
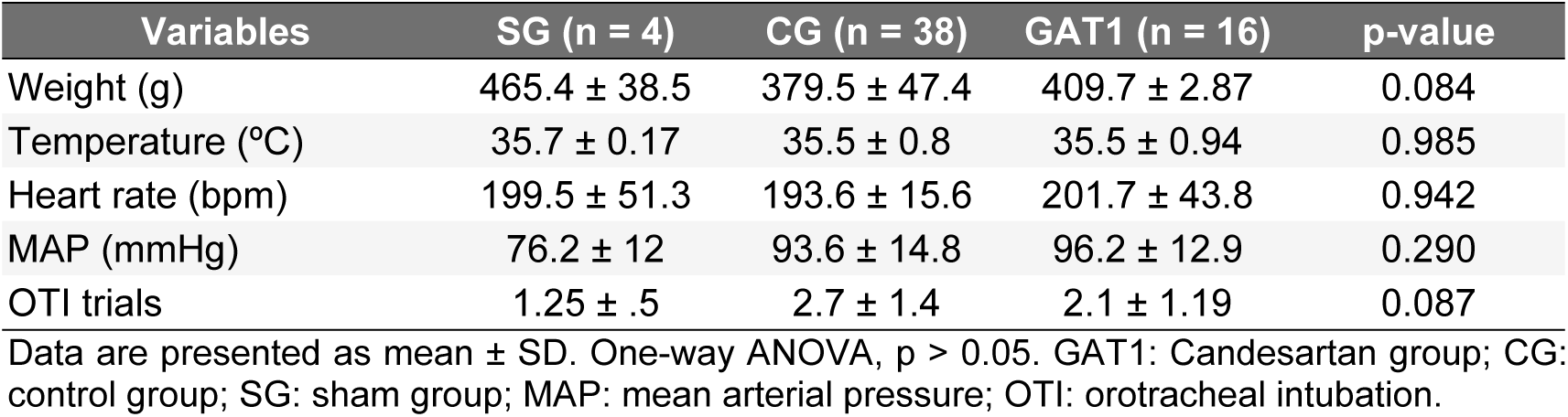
Basal characteristics of the animals.

The Candesartan group had a ROSC rate of 62.5% (10/16), higher than the control group (42.1% (16/38)) and significantly different (X^2^: 42.9; Cramer’s V: 1 .0, p = 0.00). Five animals in the control group had a non-shockable rhythm after starting the CPR maneuvers and were not submitted to a cardioverter defibrillator. Three animals in the GAT1 group had a non-shockable rhythm (p= 0.94).

All animals in the Candesartan group that met ROSC criteria survived until the end of the study. Ten (62.5%) among the 16 animals in the control group survived until the end of the established period (Figure 3) (X^2^: 4 .87, Cramer’s V: 0.487, p = 0.027).

**Fig. 3.**
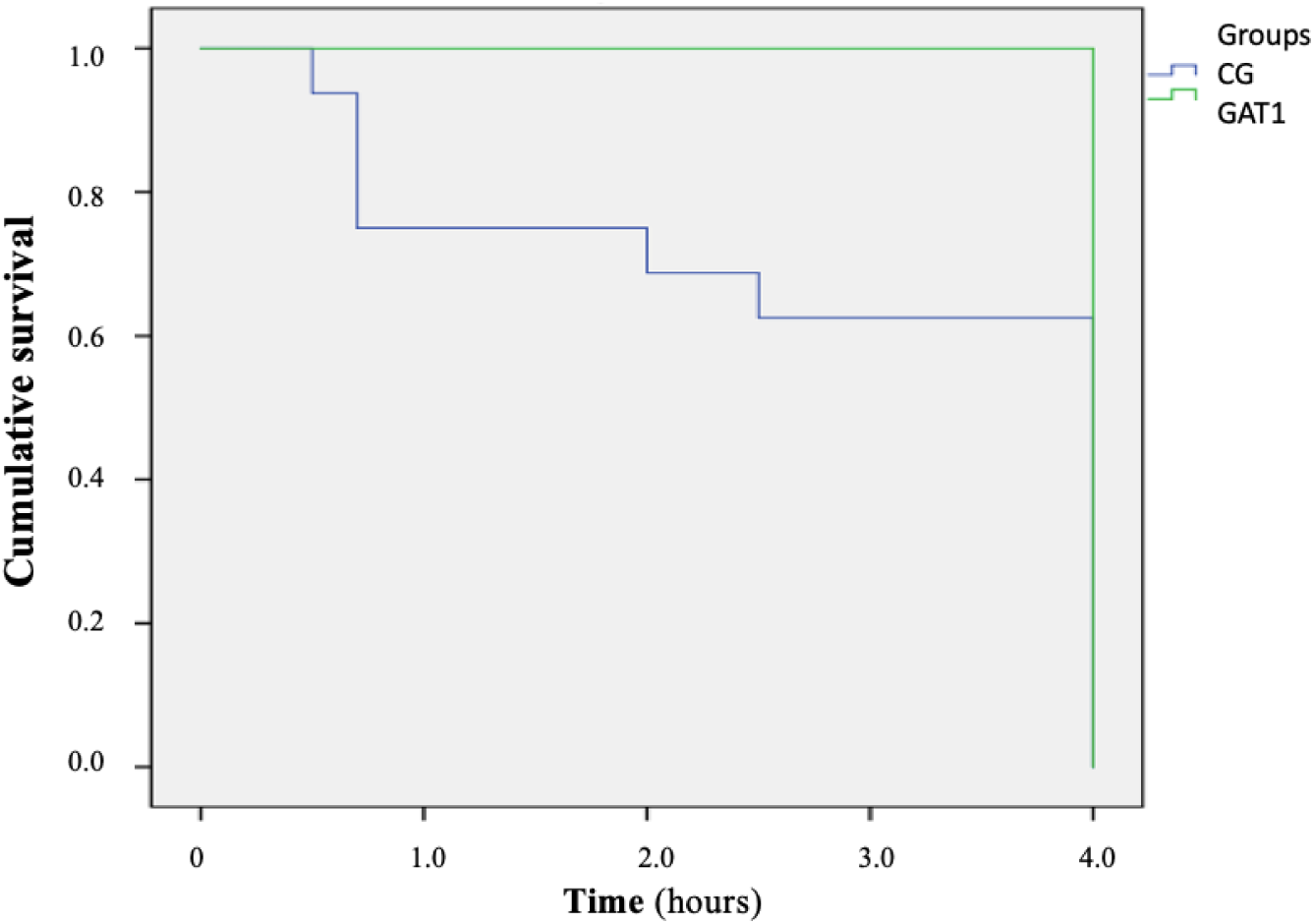
Kaplan-Meier curve of cumulative survival. There was a statistically significant difference between GAT1 and CG groups, p< 0.05

There was no difference between the means and medians of CPR duration, the number of adrenaline doses, or defibrillations (Table 2).

**Table 2.**
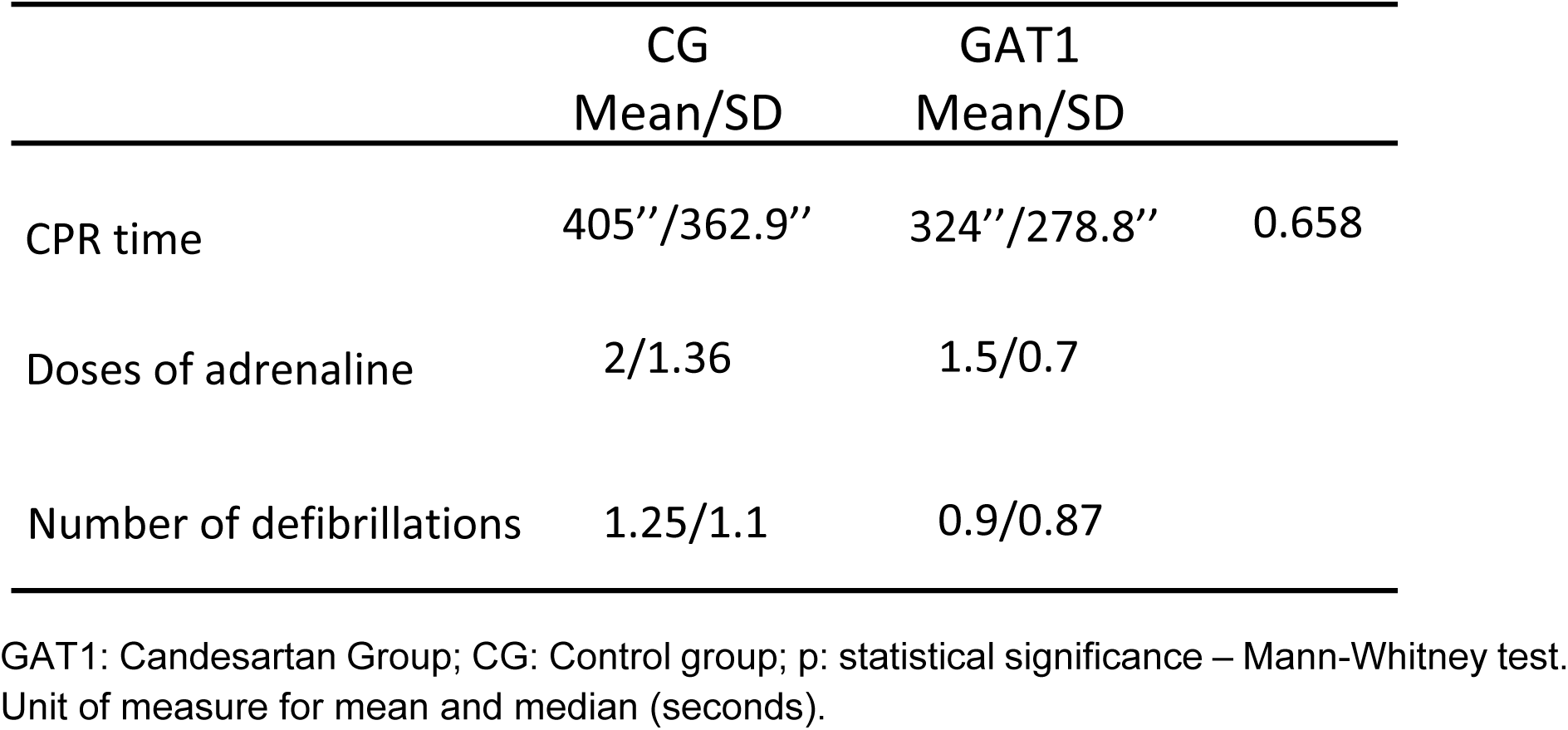
Intervention quantification during CPR.

|  | CG<br>Mean/SD | GAT1<br>Mean/SD |  |
| --- | --- | --- | --- |
| CPR time | 405''/362.9'' | 324''/278.8'' | 0.658 |
| Doses of adrenaline | 2/1.36 | 1.5/0.7 |  |
| Number of defibrillations | 1.25/1.1 | 0.9/0.87 |  |

GAT1: Candesartan Group; CG: Control group; p: statistical significance – Mann-Whitney test. Unit of measure for mean and median (seconds).

### Hemodynamic and laboratory parameters

Significant lower blood pressure in the GAT1 group was observed, starting at 40 minutes, for the animals surviving during the four hours experimental period. The same group had a lower median arterial pressure over time compared to the CG and Sham groups. The difference was statistically significant (U: 3795; p=0.041; Figure 4).

**Fig. 4.**
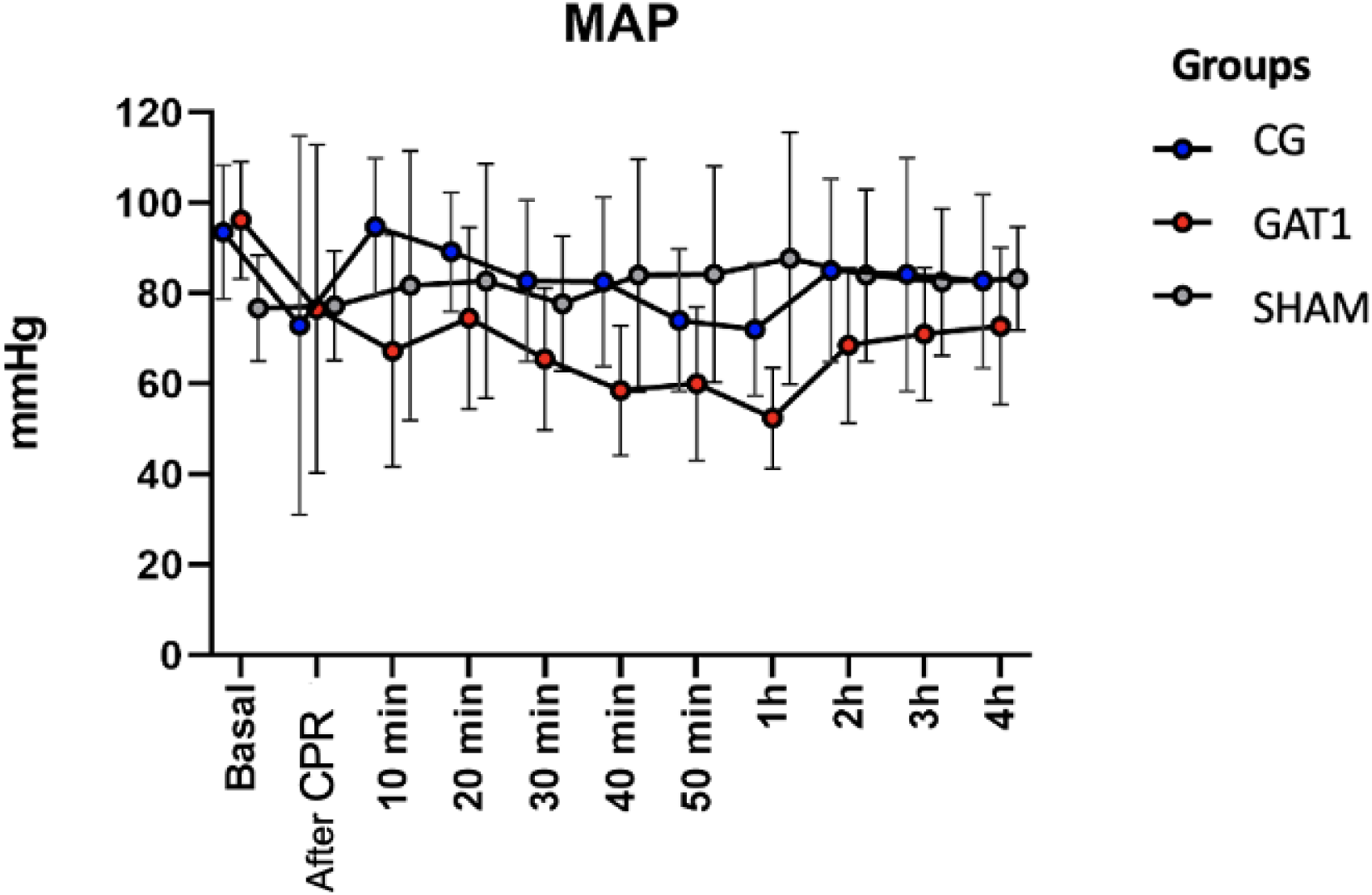
Median arterial pressure for animals that survived during the experimental period. GAT1: Candesartan Group; CG: Control group and Sham group. All met the CPR criteria.

The animals in the CG group had more cardiac arrhythmias in the ten-minute interval after ROSC. Eleven animals had disorganized heart rhythms, ten from the CG group. In these, non-sustained ventricular tachycardia (NSVT) was observed in seven animals; three had supraventricular arrhythmia. One animal from the GAT1 group had NSVT. This data was statistically significant between groups (X2: 4.85, Cramer’s V: 0.87, P = 0.000). The animals in the Sham group did not have arrhythmias during the observation period. Analyzing the temporal heart rate, it was observed that the GAT1 group had lower median HR than the CG and Sham groups, statistically significant only between 30 and 50 minutes (p= 0.037; 0.045; 0.044; Figure 5).

**Fig. 5.**
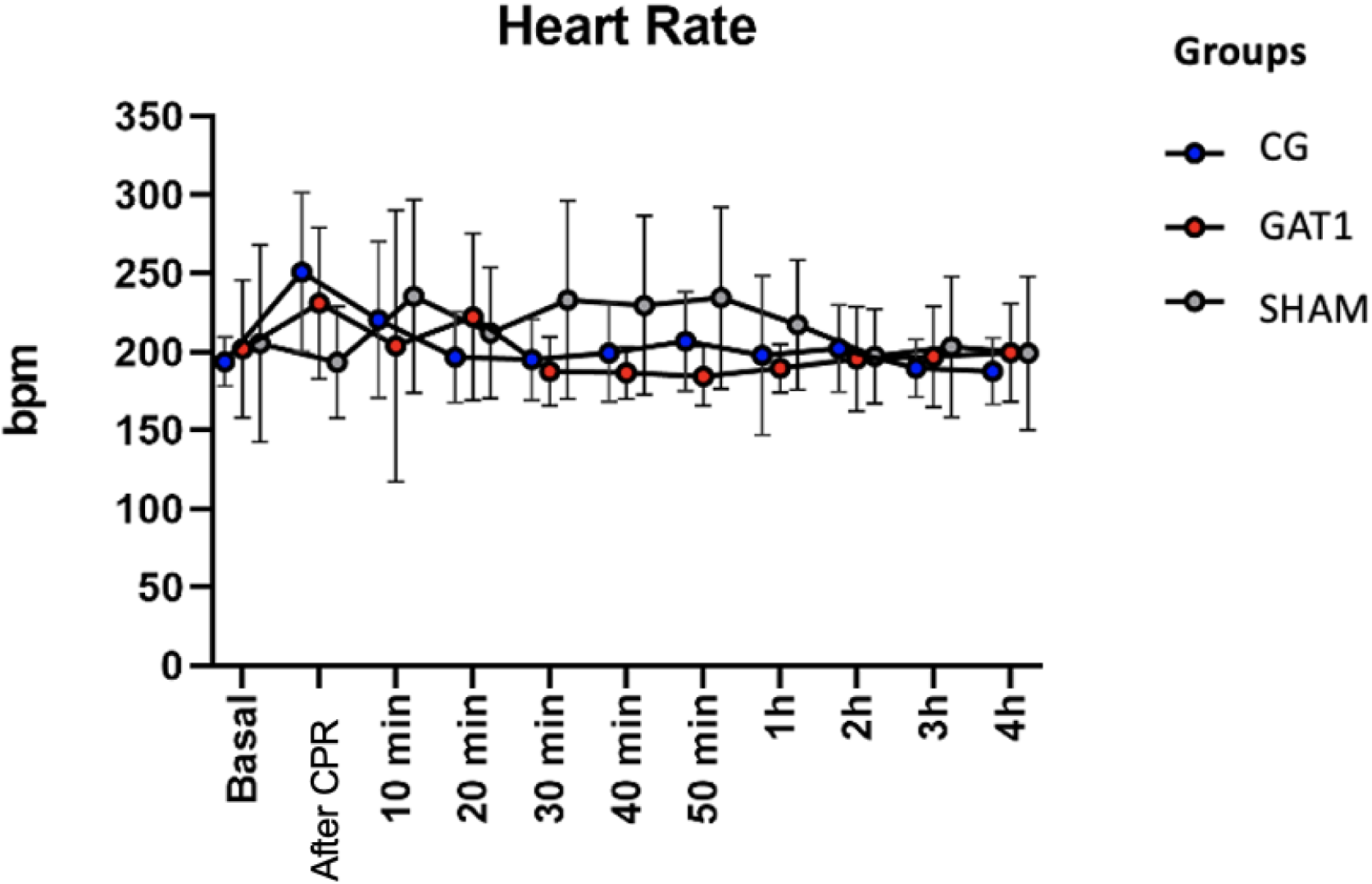
Median heart rate (bpm) among the groups of animals that survived the 4 hours of observation. GAT1: Candesartan Group; CG: Control group and Sham group. Kruskal-Wallis test.

The measurement of troponin I was performed 4 hours after animal resuscitation. CG group mean was 18265 ± 8725 pg/mL, while the GAT1 group mean was 18027 ± 8658 pg/mL, which is not statistically significant. Compared to the Sham group, these values were markedly higher (824 ± 1149 pg/mL; p=0.006).

The animals that met the ROSC criteria showed metabolic acidosis in both groups undergoing CPR. However, the GAT1 group had a higher mean pH than the control group (7.183; 7.101, U: 432, p =0.032). Similarly, higher values were observed for potassium (3.1; 3.35; U: 456 p=0.045), base excess (-9.75; -12.1; U: 434 p=0.048), and bicarbonate (17.4; 15.95, U:509, p=0.037). Data from arterial blood gas at the end of the study protocol was similar between the variables, and acidosis regulation was observed, moving towards standard values over time. Only the hematocrit of the Sham group was lower than the other groups. Medians for lactate, base excess, and bicarbonate showed no statistical difference between the groups (Table 3).

**Table 3.** Descriptive blood gas measurements in the animals that completed the study protocol.

| Variable | Time | GAT1 group | CG group | Sham group | P level |
| --- | --- | --- | --- | --- | --- |
| N | Basal | 16 | 38 | 4 |  |
|  | 10min ROSC | 10 | 16 | - |  |
|  | 4h ROSC | 10 | 10 | 4 |  |
| Temperature(°C) | Basal | 35.5±0.9 | 35.5±0.8 | 35.7±0.2 | 0.985 |
|  | 10min ROSC | 35.1±1.3 | 34.7±1.02 | 35.6±0.3 | 0.690 |
|  | 4h ROSC | 35.8±0.9 | 36.5±0.4 | 36.1±0.2 | 0.084 |
| pH | Basal | 7.41±0.04 | 7.42±0.06 | 7.36±0.07 | 0.725 |
|  | 10min ROSC | 7.170±0.103 | 7.060±0.186 |  | 0.032 |
|  | 4h ROSC | 7.358±0.101 | 7.268±0.220 | 7.310±0.070 | 0.726 |
| pCO<sub>2</sub> (mmHg) | Basal | 29.4±4.4 | 32.6±4.0 | 34.4±10.2 | 0.093 |
|  | 10min ROSC | 45.4±8.5 | 43.9±45.7 |  | 0.930 |
|  | 4h ROSC | 33.7±11.4 | 32.6±16.0 | 47.8±11.6 | 0.153 |
| pO<sub>2</sub> (mmHg) | Basal | 275.5±44.9 | 218.5±92.6 | 330.2±22.6 | 0.242 |
|  | 10min ROSC | 182.4±81.8 | 201.2±100.3 |  | 0.630 |
|  | 4h ROSC | 273.0±107.9 | 325.7±62.2 | 271.5±27.3 | 0.285 |
| Na (mmol/L) | Basal | 139.5±3.81 | 138.6±4.4 | 144.5±2.1 | 0.453 |
|  | 10min ROSC | 143.3±8.5 | 139.3±7.7 |  | 0.100 |
|  | 4h ROSC | 144.4±3.6 | 141.0±5.9 | 145.0±3.4 | 0.138 |
| K (mmol/L) | Basal | 3.6±0.4 | 4.1±0.6 | 4.0±0.3 | 0.130 |
|  | 10min ROSC | 3.1±0.5 | 3.6±3.4 |  | 0.045 |
|  | 4h ROSC | 3.9±0.6 | 4.6±1.2 | 4.7±0.7 | 0.180 |
| Cl (mmol/L) | Basal | 110.1±4.3 | 111.1±2.9 | 110.1±2.3 | 0.631 |
|  | 10min ROSC | 110.6±4.1 | 109.4±4.4 |  | 0.413 |
|  | 4h ROSC | 113.6±3.6 | 113.5±7.38 | 108.2±7.1 | 0.372 |
| Ca (mmol/L) | Basal | 0.80±0.19 | 0.91±0.18 | 0.86±0.21 | 0.216 |
|  | 10min ROSC | 0.75±0.18 | 0.89±0.19 |  | 0.037 |
|  | 4h ROSC | 0.71±0.08 | 0.89±0.22 | 0.97±0.34 | 0.112 |
| Glucose (mg/dL) | Basal | 260.0±65.1 | 295.5±75.5 | 314.0±85.6 | 0.346 |
|  | 10min ROSC | 393.0±73.6 | 441.2±106.8 |  | 0.109 |
|  | 4h ROSC | 233±88 | 234±80 | 230±139 | 0.832 |
| Lactate (mmol/L) | Basal | 3.15±1.10 | 2.79±0.95 | 2.50±0.81 | 0.320 |
|  | 10min ROSC | 7.79±3.87 | 10.64±5.93 |  | 0.257 |
|  | 4h ROSC | 2.93±1.60 | 5.78±3.89 | 2.92±1.73 | 0.341 |
| Hb (g/dL) | Basal | 16.1±1.31 | 16.7±0.92 | 16.1±2.35 | 0.356 |
|  | 10min ROSC | 15.8±1.26 | 16.6±1.53 |  | 0.215 |
|  | 4h ROSC | 16.9±1.34 | 17.4±2.05 | 15.85±1.83 | 0.350 |
| Ht (%) | Basal | 49.2±3.97 | 49.2±5.85 | 46.3±4.04 | 0.235 |
|  | 10min ROSC | 48.6±3.8 | 51.3±5.3 |  | 0.206 |
|  | 4h ROSC | 51.9±4.0 | 53.1±6.1 | 45.8±3.2 | 0.037 |
| BE (mmol/L) | Basal | -2.63±1.5 | -4.11±1.6 | -2.87±1.2 | 0.406 |
|  | 10min ROSC | -10.6±4.9 | -14.9±8.0 |  | 0.048 |
|  | 4h ROSC | -5.9±5.3 | -9.1±10.4 | -2.5±3.7 | 0.342 |
| Bic (mmol/L) | Basal | 19.6±1.6 | 21.3±1.5 | 21.6±2.3 | 0.056 |
|  | 10min ROSC | 16.6±3.7 | 13.5±6.3 |  | 0.037 |
|  | 4h ROSC | 18.7±4.8 | 16.7±8.2 | 23.2±3.7 | 0.180 |

Descriptive measures of arterial blood gas baseline parameters after the return of spontaneous circulation and at the end of the study. Kruskal-wallis test. BE: base excess; Bic: bicarbonate.

### Cerebral and myocardial injury

Microscopic evaluation showed more red neurons and signs of necrosis throughout the parietal cortex and hippocampus in the CG group compared to the GAT1 group (p= 0.025, Cramer’s V 0.539). The Sham group showed no signs of neuronal injury (Figure 6A). The distribution of neuronal injury scores between the groups is seen in Figure 7.

**Fig. 6.**
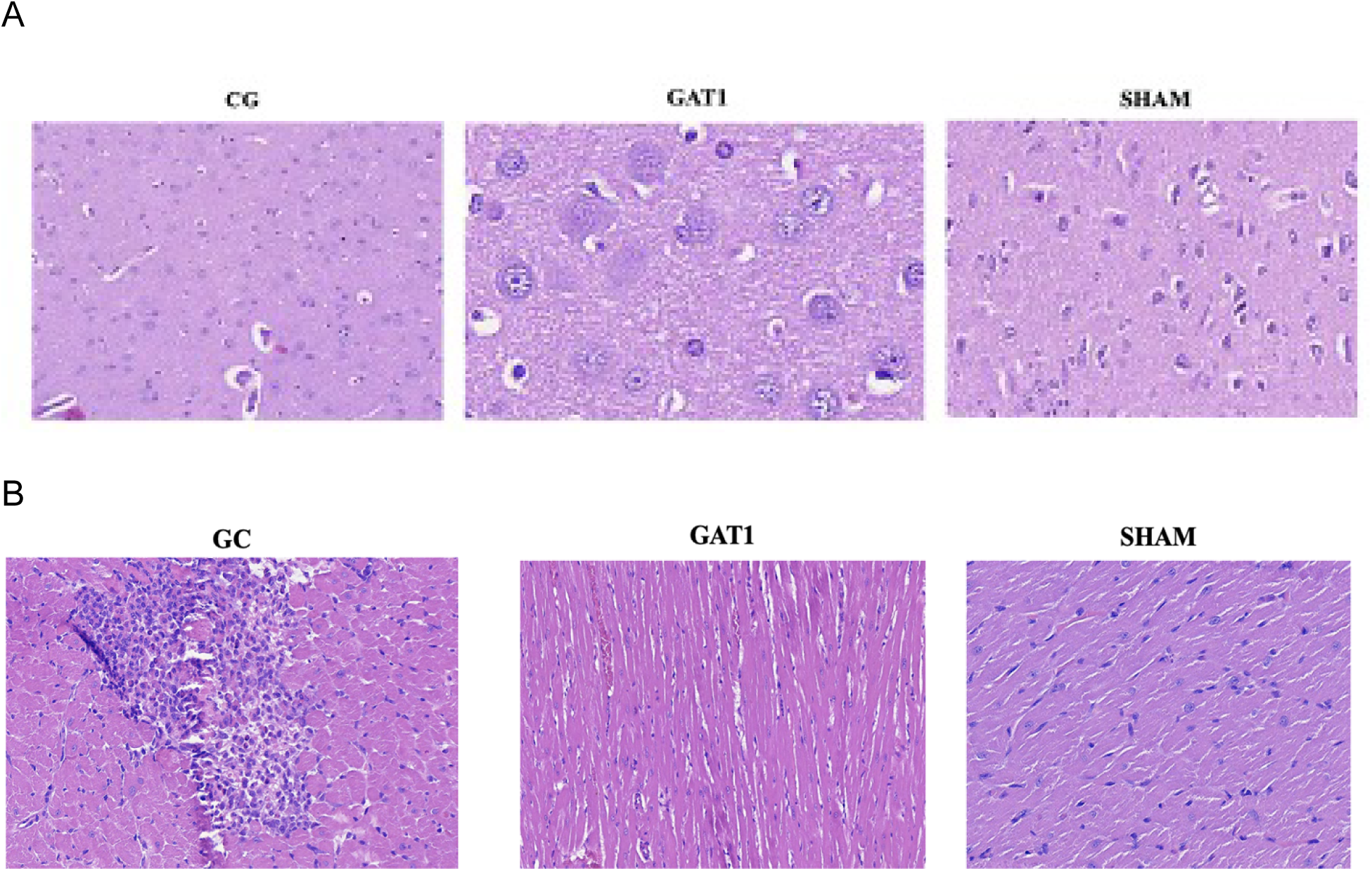
A. Micrographs of brain tissue, stained with hematoxylin and eosin, 40x magnification, blue arrow indicates red neuron. B. Micrographs of the myocardial tissue stained with hematoxylin and eosin (10 fields/animal).

**Fig. 7.**
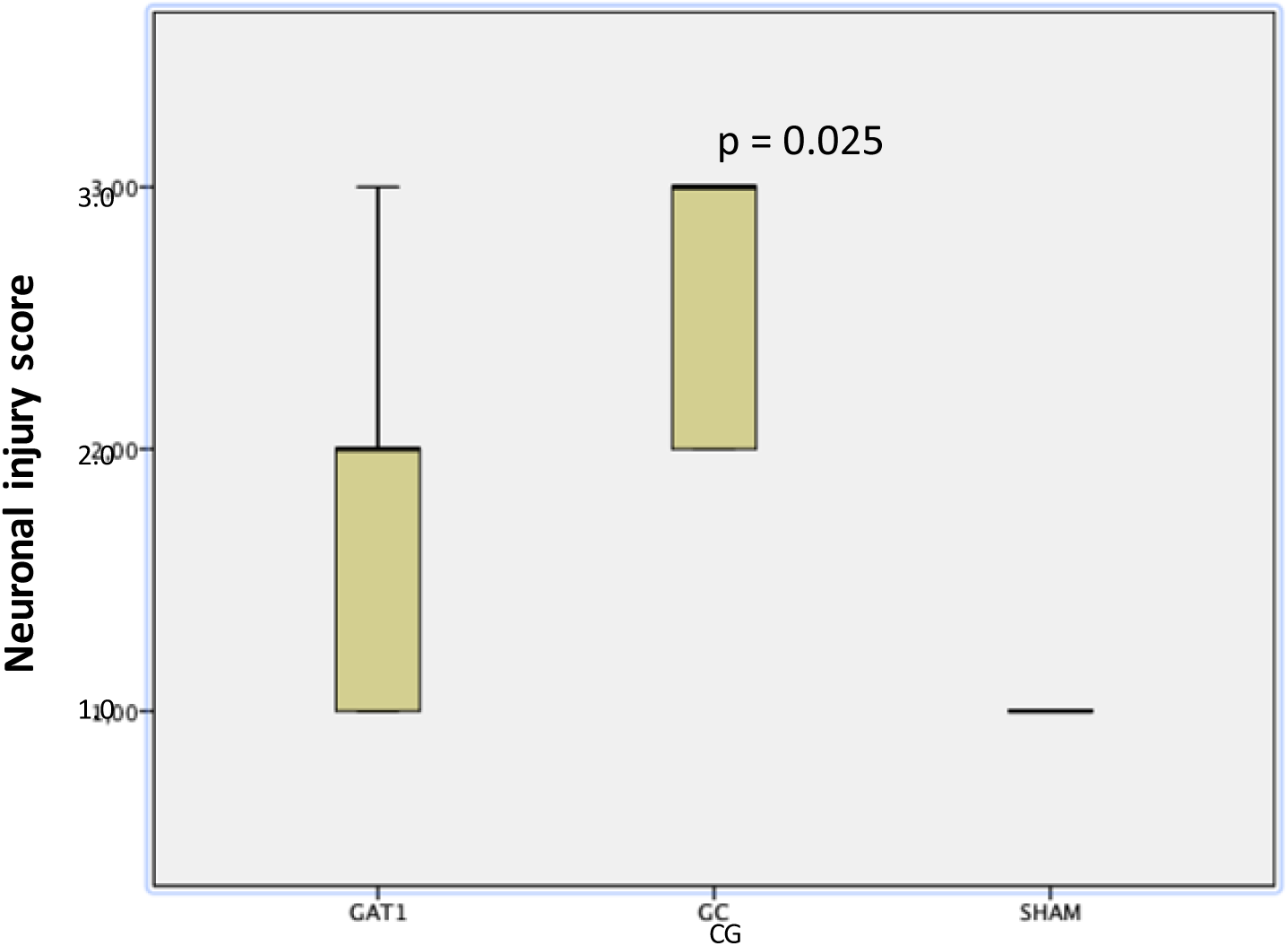
Neuronal injury score after four hours of observation in animals meeting the ROSC criteria. GAT1: angiotensin 1 receptor antagonist group, ROSC: return of spontaneous circulation. Qui square Test.

Microscopic evaluation showed that myocardial cells were orderly arranged with intact nuclei in the Sham group. In the CG group, myocardial cells were irregular, and many immune cells were aggregated (Figure 6B). Compared to the GAT1 group, fewer myocardial cells were disorganized, and cell infiltrations were lower in the latter group (Figure 6B). The CG’s myocardial damage score increased significantly compared with the Sham group (Figure 8). Furthermore, the myocardial damage score significantly decreased in the GAT1 group compared to the CG group, as seen in Figure 8.

**Fig. 8.**
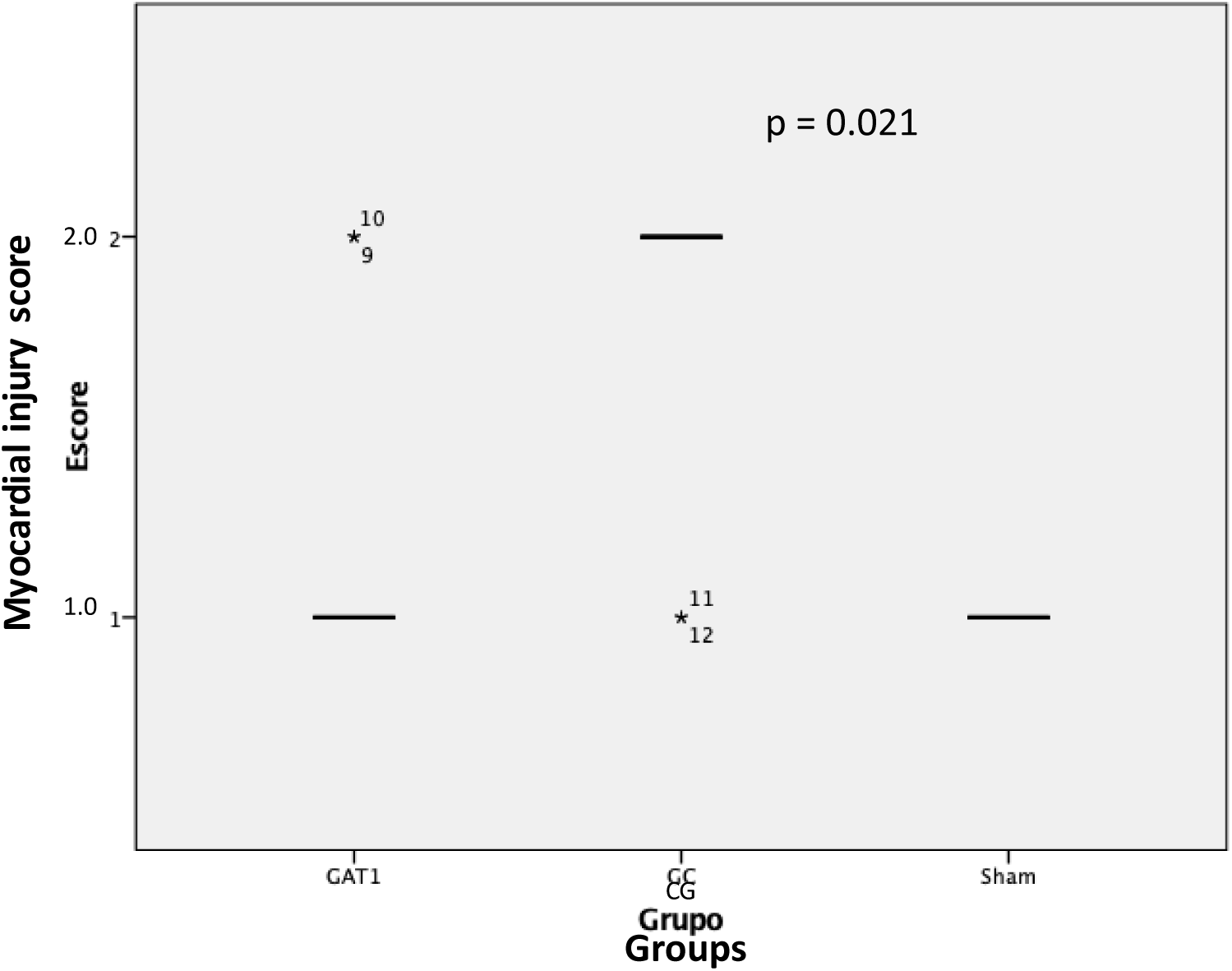
Myocardial injury score after four hours of observation in animals meeting the ROSC criterion. GAT1: angiotensin 1 receptor antagonist group, CG: control group, ROSC: return of spontaneous circulation. Qui square Test.

When using the TUNEL method, similar medians were observed for the number of apoptotic cells in both brain and heart groups. The Sham group showed no signs of cell injury.

## Discussion

This is the first report evaluating the effect of AT1 receptor blockade during cardiac arrest as an adjunct to advanced life support. We demonstrated that blocking the Angiotensin II AT1 receptor using candesartan leads to a higher rate of ROSC and survival in rats after CPR. The results indicate that candesartan attenuates neuronal and myocardial injury when administered during CPR maneuvers.

Previous experimental studies have evaluated the role of this receptor blockade under focal I/R injury, submitting the animals to a regimen of drug therapy prior to the CPR event or using models of isolated hearts following the Langendorf technique ^15, 17–21^. Furthermore, most of these studies did not use angiotensin in the perfusate, precluding any conclusion about the effects of inhibiting the receptor of this enzyme. Since the renin-angiotensin system is related to worsening reperfusion injury, adding angiotensin to the perfusate is necessary as a model for investigating pharmacological approaches that improve cardiac function after ischemia-reperfusion ^22, 23^.

Currently, most studies optimize brain protection after I/R episodes. Regarding cardioprotective strategies after ROSC, published data are scarce. Therefore, finding interventions that effectively improve post-resuscitation myocardial and neuronal dysfunction is critical for patient prognosis.

The findings from brain histopathology point to candesartan as a neuroprotector in ischemic injury caused by CPR. This is due to the blockade of angiotensin II, the main effector/ligand of the brain’s intrinsic renin-angiotensin system. It is a neuropeptide produced mainly by glial cells in the central nervous system. Hyperactivation of the brain AT1 receptor is responsible for the harmful effects associated with the RAS, leading to vasoconstriction, cerebral blood flow decrease, increased oxidative stress, and vulnerability to ischemia, in addition to promoting vascular and tissue inflammation and neurodegeneration exacerbation^24^. AT2 stimulation counteracts these mechanisms. Therefore, selective blocking of AT1 receptors with Angiotensin Block Receptors (ABRs), especially candesartan, which crosses the blood-brain barrier ^25^, may offer superior protection than simultaneously decreasing AT1 and AT2 receptors, such as angiotensin-converting enzyme inhibitors (ACE inhibitors). Such mechanisms were used by Hajjar et al. (2020) ^26^, demonstrating less neurocognitive impairment in an elderly population that used candesartan instead of Lisinopril, regardless of blood pressure control.

Cardiac dysfunction associated with post-CPR syndrome is characterized by severe myocardial impairment and global hypokinesia, affecting the success rate of cardiopulmonary resuscitation. Cardiac function deterioration begins minutes after arrest and peaks within 2 to 5 hours after resuscitation, reinforcing early intervention and treatment ^27^. Proposed molecular mechanisms for post-resuscitation myocardial dysfunction involve I/R injury, resulting in large amounts of oxygen free radicals that damage cell membranes and induce myocyte necrosis and apoptosis. Moreover, an intracellular accumulation of Na+ is caused by the cytosolic overload of Ca^2+^ through the action of the Na^+^/Ca^2+^ exchanger present in the sarcolemma membrane ^28^.

The presence of endogenous or paracrine RAS in the heart has been recently recognized. RAS components – angiotensinogen and renin messenger RNA, angiotensin I to angiotensin II (Ang II) converting enzyme – have previously been detected in the heart and seem functionally integrated ^11–14^. Cardiac Ang II may be involved in regulating coronary blood flow, modulation of sympathetic neurotransmission, cardiac contractility, and stimulation of cell hyperplasia and hypertrophy, in addition to repairing the cardiovascular structure. Angiotensin II interacts with two pharmacologically distinct subtypes of cell surface receptors, type 1 and type 2 Ang II receptors. Type 1 receptors mediate the main cardiovascular effects of Ang II ^29^. Such mechanisms may explain the significant differences found in the lower myocardial injury score in the GAT1 group reported here.

Furthermore, limiting Na^+^ influx into the sarcolemma during ventricular fibrillation resuscitation prevents the accumulation of excess mitochondrial Ca^2+^ and attenuates myocardial injury ^30^. This mechanism may explain the significant reduction in cardiac arrhythmias observed here since no differences in CPR duration, number of defibrillations, or adrenaline doses between groups were observed. Thus, we suggest that the action of candesartan does not interfere with the reversal of heart rhythm (VF or VT) during CPR. However, as the incidence of cardiac arrhythmias one hour after ROSC was higher in the control group, the drug could potentially affect the reduction of new arrhythmogenic triggers. This mechanism leads to the blockade of Angiotensin II AT1 receptors, explaining the higher survival of the animals in the GAT1 group. This mechanism has already been demonstrated in previous studies ^12, 14^.

As for the metabolic analysis of I/R, the GAT1 group had a lower level of acidosis and base intake than the CG group, suggesting that the combination of adrenaline and candesartan can protect myocardial tissue and improve energy metabolism after ROSC. The best survival rate for the GAT1 group corroborates this hypothesis. Furthermore, the lower blood pressure in the GAT1 group compared to the control and Sham groups may be due to the hypotensive effect of angiotensin II antagonism ^10^. This finding may compromise the translation of this experimental study into clinical studies since we do not know the behavior of this drug after CPR in humans.

### Limitations

The main limitations of this study include the absence of a group that received candesartan alone during resuscitation maneuvers. However, the use of adrenaline as a significant factor for ROSC increase in rats has already been documented ^15^. Another limitation was that the ROSC rate was lower than the experimental model adopted by our research group ^15^, requiring more animals to reach 10 per intervention and control group, as indicated in the study design. Although based on estimated sample calculation, the small sample size may reduce the study’s statistical power.

Our results suggest excellent perspectives for using angiotensin II AT1 receptor antagonists for cardiopulmonary resuscitation protocols, especially in assisted arrests, where the intervention time is shorter compared to out-of-hospital CAs.

## Authors contribution

Araújo Filho, EAF: methodology, writing, experimentation, article writing, and final draft approval; Carmona, MJC: methodology preparation, last draft revision; Otsuki, DA: surgical instrumentation of the animals and data collection, literature review, and statistical analysis; Maia, DRR: data collection, review of statistical analysis, and laboratory tests; Lima, LGCA: histopathological analysis of the brain and myocardial tissues of the animals; Vane, MF5: project preparation, coordinator of the work execution, final draft revision and statistical analysis.

## Acknowledgments

We thank the staff and members of the Anesthesiology Department from Faculdade de Medicina da USP for their support.

## Conflict of interest

The authors declare no conflict of interest.

## Funding

There was no funding for the work.

---

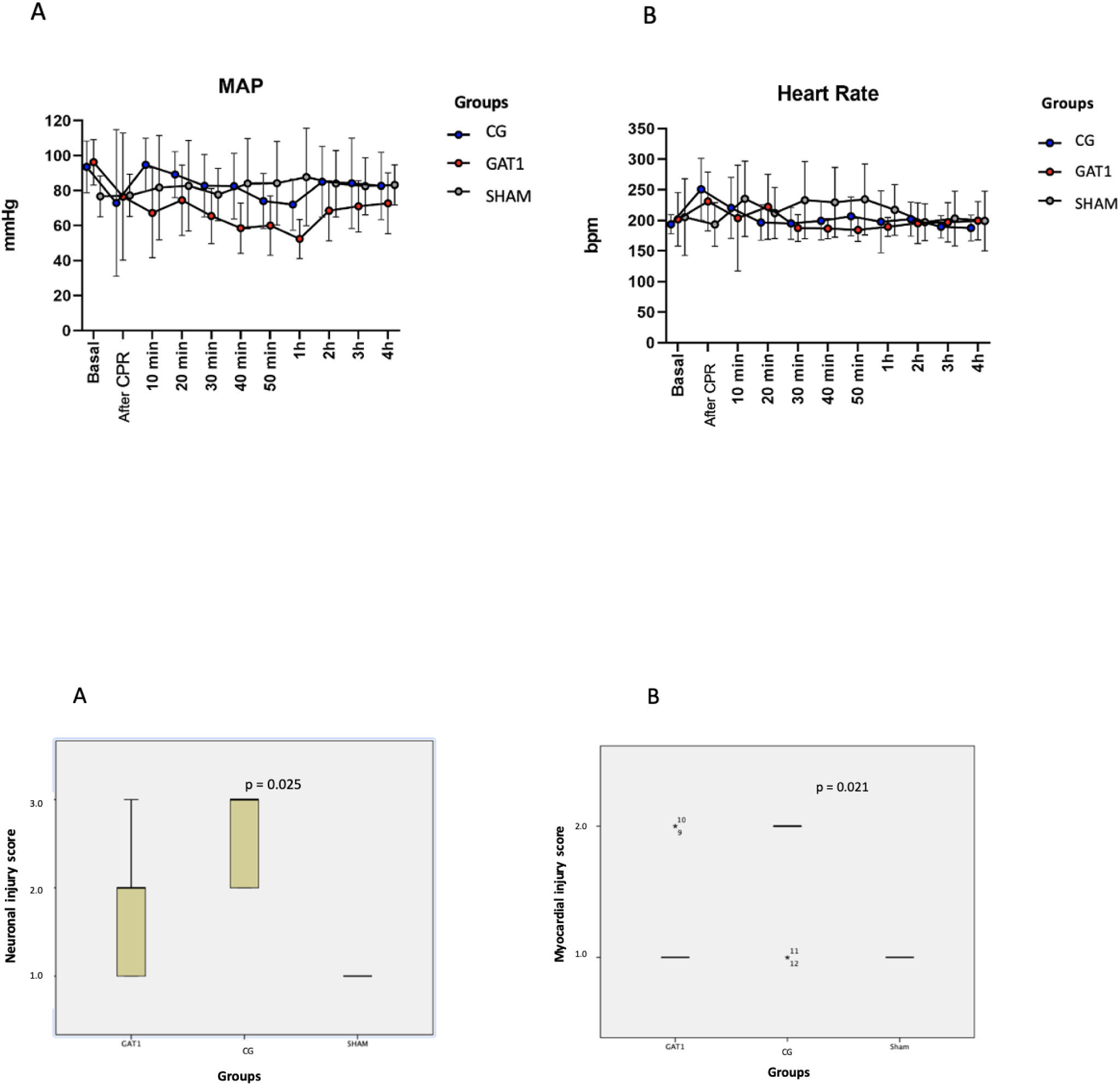

## Notes

### Competing Interest Statement

The authors have declared no competing interest.

